# Integrative Modeling of a Sin3/HDAC Complex Sub-structure

**DOI:** 10.1101/810911

**Authors:** Charles A. S. Banks, Ying Zhang, Sayem Miah, Yan Hao, Mark K. Adams, Zhihui Wen, Janet L. Thornton, Laurence Florens, Michael P. Washburn

## Abstract

Sin3/HDAC complexes function by deacetylating histones, which makes chromatin more compact and modulates gene expression. Although components used to build these complexes have been well defined, we still have only a limited understanding of the structure of the Sin3/HDAC subunits as they are assembled around the scaffolding protein SIN3A. To characterize the spatial arrangement of Sin3 subunits, we combined Halo affinity capture, chemical cross-linking and high-resolution mass spectrometry (XL-MS) to determine intersubunit distance constraints, identifying 66 high-confidence interprotein and 63 high-confidence self cross-links for 13 Sin3 subunits. To validate our XL-MS data, we first mapped self cross-links onto existing structures to verify that cross-link distances were consistent with cross-linker length and subsequently deleted crosslink hotspot regions within the SIN3A scaffolding protein which then failed to capture crosslinked partners. Having assessed cross-link authenticity, we next used distance restraints from interprotein cross-links to guide assembly of a Sin3 complex substructure. We identified the relative positions of subunits SAP30L, HDAC1, SUDS3, HDAC2, and ING1 around the SIN3A scaffold. The architecture of this subassembly suggests that multiple factors have space to assemble to collectively influence the behavior of the catalytic subunit HDAC1.

## Introduction

Although solution NMR and crystallographic studies have provided valuable insight into the structure of components of macromolecular complexes, it is often challenging to determine the architecture of subunits as they are assembled into higher-order structures. Crystallographic studies are limited by the requirement that the molecules isolated can form rigid crystals suitable for structure determination^1^. In addition, NMR studies of larger proteins and protein complexes are hindered by the large number of NMR signals generated which cause spectral crowding^2^. Recent developments in cross-linking techniques combined with advances in high-resolution mass spectrometry have provided valuable new tools to address these limitations^3^.

Pioneering work by Rappsilber et al. in 2000^4^ established protein complex organization by combining cross-linking with mass spectrometry to investigate the spatial organization of the yeast nuclear pore complex. They isolated populations of cross-linked Nup84p complex subunits using SDS-PAGE gels and excised 8 bands corresponding to groups of 2-5 crosslinked proteins for identification by MALDI-MS. Using this approach, they determined the relative positions of the subunits in the complex. Later, in 2007, Maiolica et al. combined this approach with improved database searching techniques to identify the location of specific cross-links, mapping 26 cross-links between four subunits of the NDC80 complex^4^. More recently, Kao et al. developed an MS cleavable crosslinker, DSSO, which can be combined with high-resolution mass spectrometry to improve unambiguous identification of cross-linked peptides^5,6^.

In this study, we combine this approach with Halo affinity purification^7^ to capture positional information for Sin3 complex subunits in solution. High-confidence residue-specific interactions are identified by integrating MS1, MS2 and MS3 data analyses. Following protein complex isolation, cross-linking, tryptic digestion, and reverse phase peptide separation, putative crosslinked peptides with high charge (z ≥ +4) can be isolated for further analysis following MS1. MS2 analysis confirms that a cross-linked peptide has been isolated. During MS2, cross-links between linked peptides are cleaved using CID (collision induced dissociation) resulting in a characteristic pattern of two pairs of ions in the MS2 spectra characteristic of a cross-linked peptide. As CID can break either of two bonds within DSSO, a separated peptide contains a lysine modified by either a thiol or alkene group originating from the cross-link. The ~32 Da mass difference of these alternate modifications results in the characteristic pair of ions corresponding to each separated peptide. High-resolution mass spectrometry enables subsequent MS3 sequencing of these four fragments using higher energy CID. Thus, high confidence, low abundance cross-links can be rapidly identified. Here, we combine this approach with Halo affinity purification to investigate the subunit architecture of histone deacetylase complexes.

Sin3/HDAC complexes influence transcriptional control on subsets of genes by modulating the chromatin environment. They function by orchestrating the deacetylation of lysine residues on N-terminal histone tails using the catalytic subunits HDAC1 and HDAC2. This results in chromatin compaction and the subsequent repression of gene transcription as genes become inaccessible to the transcriptional machinery. The precise targeting of gene repression by Sin/HDAC mediated histone deacetylation is likely orchestrated by the non-catalytic Sin3 subunits^8^, since HDAC1/2 are not unique to Sin3 and are used by other histone deacetylase complexes, including NuRD^9^ and CoREST^10^. Although the subunit composition of Sin3/HDAC complexes has been established^11^, how they organize around SIN3A to accomplish HDAC1/2 mediated deacetylation of specific histone residues at specific genomic loci remains unclear.

Uncovering the architecture of Sin3/HDAC complexes is essential in understanding the contribution of individual subunits to their function which in turn is vital in understanding how misregulated Sin3 complexes contribute to human disease. SIN3A, the scaffolding protein around which the complex assembles, is frequently mutated in human cancers^12^, and Sin3 complexes offer likely therapeutic targets for a variety of diseases^12^ including triple negative breast cancer^13^ and pancreatic cancer^14^. Current therapeutic strategies using HDAC inhibitors such as vorinostat (suberanilohydroxamic acid or SAHA) are not specific, targeting a variety of HDAC containing complexes^15^. Targeting HDAC activity within the context of Sin3 complexes more specifically will require a more sophisticated understanding of how Sin3 subunits cooperatively control HDAC1/2 recruitment and function.

Here, we isolated Sin3/HDAC complexes using a Halo-tagged version of the SAP30L subunit stably expressed in Flp-In™-293 cells and captured positional information for individual Sin3 subunit residues using the cross-linker DSSO. Following high-resolution mass spectrometry, we identified 63 Sin3 subunit self cross-links and 66 Sin3 subunit interprotein crosslinks. We next used previously determined structures to confirm that the distances observed between cross-linked subunits were consistent with the distance limits required by the ~10 Å DSSO cross-linker. We further judged the validity of our cross-linking data by determining whether SIN3A crosslink hotspots were indeed required for capturing cross-linked subunits. Finally, we used intersubunit cross-links to dock SAP30L, SIN3A, and HDAC1 structures and to map the relative locations of SUDS3, SAP130, HDAC2, and ING1 on the resulting structure. Importantly, this reveals the position of the HDAC1 active site relative to other subunits. Using molecular modeling to integrate a comprehensive map of cross-links between Sin3 subunits with existing structural data has revealed the arrangement of subunits at the core of the Sin3 complex, illuminating how they might function collectively to regulate chromatin accessibility and gene transcription.

## Results

### Previous studies provide a framework for developing a high-resolution network of Sin3 subunit interactions

determining how groups of Sin3 complex subunits are assembled is important in understanding how they function together to control the status of histone acetylation and hence regulate gene expression. Several important studies have enabled a progressively more detailed picture of Sin3 subunit interactions to emerge (Fig. 1, Supplementary Table 2). In 1997, Laherty et al. first established that a ~375 amino acid conserved domain within the Sin3 scaffolding protein mSin3A was important for its interaction with the catalytic subunit HDAC2. They named this region the <u>H</u>DAC <u>I</u>nteraction <u>D</u>omain or HID^16^. Later studies determined further interactions between SIN3A and other subunits. Importantly, Zhang et al. used GST pulldown assays to establish direct interactions between GST-SAP30 and mSin3A and between GST-SAP30 and HDAC1^17^, suggesting that these three proteins functioned in close proximity. Later, the interactions between SIN3A, HDAC1, and SAP30 were further refined^18–21^, and interactions between the SIN3A HID and two other subunits, SUDS3 and SAP130^18,19,21^, were established. In particular, Xie et al. determined a structure of part of the C-terminus of SAP30 in complex with the PAH3 domain of SIN3A^20^, and Clark et al. determined a structure explaining an interaction between part of SUDS3 with part of the SIN3A HID^21^. Thus, the PAH3/HID region within SIN3A was established as a central organizing platform around which several other Sin3 components (SAP30, HDAC1, SUDS3, and SAP130) might assemble. Despite these advances, it remains unclear whether there is space for these components to dock together on this platform within SIN3A.

**Figure 1.**
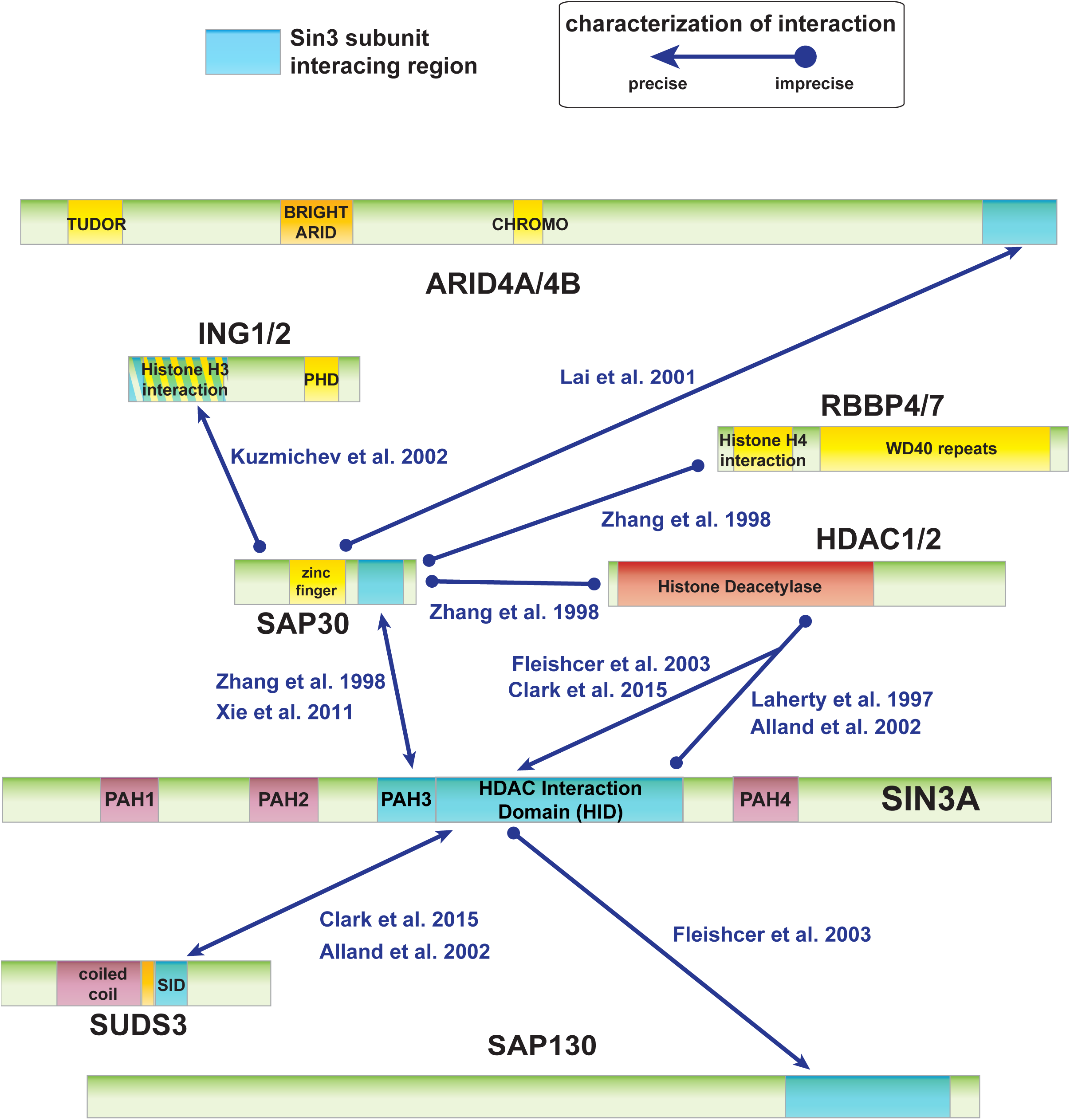
Previously characterized interactions between Sin3 subunits. A summary of the interactions characterized and details of each reference is provided in Supplementary Table 2. Regions important for interactions between Sin3 subunits are shown in blue, for interaction with DNA/nucleosomes in yellow, for histone deacetylase activity in red, and other structurally defined regions in purple.

### From AP-MS to XL-MS: mapping proximal amino acids among Sin3 complex components

To address how Sin3 subunits might be organized around the SIN3A HID, we used a cross-linking mass spectrometry (XL-MS) approach to determine proximity constraints for pairs of amino acids within Sin3 complexes. Previously, we had determined the set of Sin3 subunits copurifying with SAP30L, a SAP30 homolog, using an affinity purification mass spectrometry (AP-MS)^11^ approach (Supplementary Figure 1). To extend this analysis, we now treated purified SAP30L containing complexes with the MS cleavable crosslinker DSSO prior to liquid chromatography mass spectrometry (LC/MS) analysis, capturing additional structural information by highlighting pairs of residues within Sin3 assemblies that likely reside within a distance of < 30 Å (Fig. 2*A,B*, Supplementary Figure 2).

**Figure 2.**
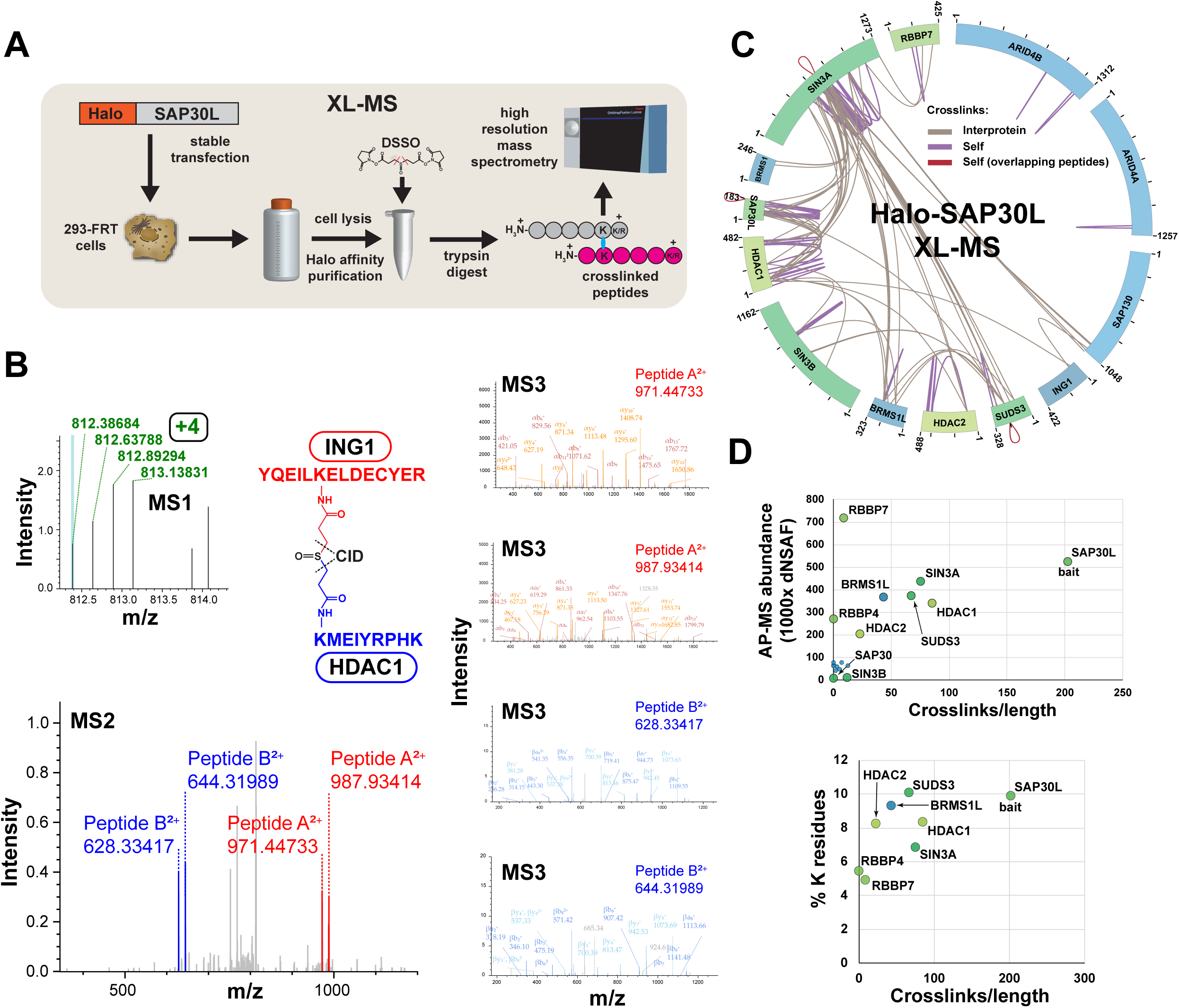
MS cross-link analysis of Sin3/HDAC complexes. (A) Workflow for XL-MS experiments. In contrast to AP-MS experiments, Halo purified samples were treated with DSSO prior to analysis by high-resolution mass spectrometry. (B) High-resolution MS2 and MS3 spectra used to identify the ING1-HDAC1 cross-linked peptide. First, putative cross-linked peptides with charge ≥ +4 were selected during MS1 analysis. Next, low energy CID was used to cleave the DSSO crosslinker, generating a pair of fragments for each peptide sequence. These were identified using MS2 (ING1 peptides shown in red, HDAC1 peptides in blue). Finally, the four fragments were sequenced using MS3. Annotated spectra were generated using Proteome Discoverer™ software version 2.1. (C) Cross-link map for the Sin3/HDAC complex. Cross-link identifications are from three XL-MS experiments. Interprotein cross-links are shown in brown, self cross-links in purple, and self cross-links with overlapping peptides in red. Values indicate protein length (amino acids). Details of cross-links are in Supplementary Table 3. (D) Relationship between observed cross-link abundance and either protein abundance or lysine content for Sin3 subunits. Crosslink abundance was calculated as 1000 x (semi-crosslinks/protein length (aa)), with two semi-cross-links counted for each of the protein’s self cross-links and one for each of the protein’s interprotein cross-links. Protein abundance dNSAF values (four biological replicates) for SIN3 subunits copurifying with Halo-SAP30L were published previously (refer to Banks et al. Supplementary Table S3^11^).

We identified 66 interprotein cross-links between different Sin3 subunits and we identified 63 self cross-links within ten Sin3 subunits (Fig. 2*C*, Supplementary Table 3). It is not possible to tell whether these self cross-links result from cross-links within a single molecule or from cross-links between two identical molecules (for example if the subunit exists as a homodimer). However, for some self cross-links within the subunits SIN3A, SAP30L, and SUDS3, the sequences of cross-linked peptides overlap. This suggests that these subunits might form homodimers (red cross-links Fig 2*C*). Indeed, Clark et al. had previously shown that SUDS3 has homodimerization activity^22^ and proposed that this might enable a pair of Sin3 complexes to assemble together, bridging two adjacent nucleosomes^21^.

The cross-links did not appear to be distributed evenly amongst the 13 Sin3 subunits (Fig. 2*C*). While some quite large proteins had few cross-links (e.g. one self cross-link detected for ARID4A), most crosslinks were distributed amongst five proteins: SIN3A, SAP30L, HDAC1, SUDS3, and BRMS1L. In addition, there appeared to be “hotspots” of cross-links within proteins, for example the HID region within SIN3A. To explain the uneven distribution of cross-links, we first assessed whether the paucity of cross-links on certain subunits might simply reflect a low abundance of these subunits in our purifications. To test this, we calculated a factor reflecting the number of cross-links per unit length of the protein (semi-crosslinks per 1000 amino acids) and compared this with protein abundances for the various Sin3 subunits derived from our AP-MS studies (Fig. 2*D*). Although low abundance can explain the deficit of crosslinks identified for some proteins (SIN3B, SAP130, ARID4A/B, SAP30, BRMS1 and FAM60A), it did not explain the deficit of cross-links for the relatively abundant RBBP4 and RBBP7 subunits. A second possibility was that the cross-link deficit for RBBP4/7 might be explained by a low number of lysine residues in these proteins. When we calculated the percentage of lysines for the eight most abundant subunits, we indeed found that RBBP4/7 have the lowest percentage of lysines, and this might partially explain their lack of cross-links. In addition to their low lysine content, RBBP4 and RBBP7 are also largely formed from beta sheets, and previous studies have proposed that these structures often correlate with low levels of cross-links^23^.

### Euclidean distances between cross-linked residues mapped to Sin3 structures are consistent with cross-linker length

To further assess our crosslinking data, we tested whether the distances between cross-linked residues, which mapped to experimentally determined Sin3 tridimensional structures, were consistent with the length of the DSSO cross-linker. Although the spacer length of DSSO is 10.1 Å, Merkley et al. had previously determined that distances of up to 30 Å between alpha carbon atoms of crosslinked residues were appropriate in their analysis of the similarly sized cross-linker DSS^24^. We first assessed 11 crosslinks that mapped within the SIN3A structure 2N2H^21^ (Fig. 3*A*) and determined that 10 of these corresponded to Cα-Cα distances of < 30 Å (Fig. 3*B*). Curiously, we found one crosslink between two residues within the structure with a much longer Cα-Cα distance of 44 Å. It is possible that either this 44 Å crosslink is between two different SIN3A molecules, or that other conformations of this region exist in solution, especially since one of the two linked lysines is located at the end of the C-terminal α-helix in the 2N2H partial structure that could be folded differently in the context of full-length SIN3A and the assembled SIN3 complex. In addition to the SIN3A structure 2N2H, we found four other structures mapping to regions of other Sin3 subunits containing self cross-links. The structures 2N1U^25^ and 2LD7^20^ both map to SAP30L, the structure 5ICN^26^ maps to HDAC1, and the structure 3CFV^27^ maps to RBBP7 (Fig. 3*C*). Of the 23 cross-links that mapped within existing structures, 22 (96%) had corresponding Cα-Cα distances of < 30 Å, confirming that the self cross-links that we identified likely originate from intact Sin3 structures.

**Figure 3.**
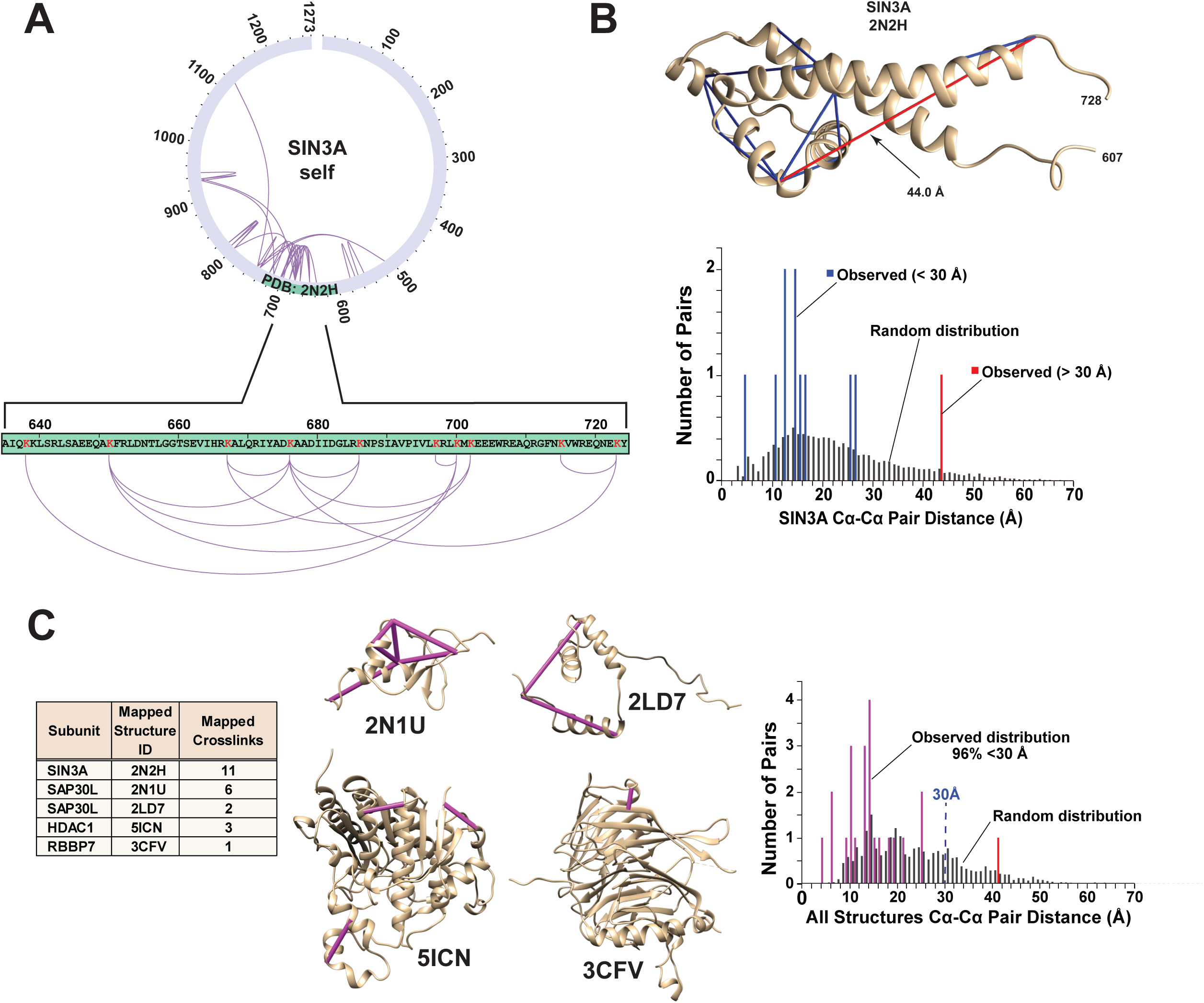
Cα–Cα distance distributions for cross-links mapping to Sin3 subunit structures. (A) The structure PDB: 2N2H^21^ maps to a SIN3A region containing 11 self cross-links. (B) Distribution of Cα–Cα crosslink distances mapping to 2N2H (experimentally determined distances (blue/red bars) and random probability distribution (grey bars)). Distances were measured xiVIEW^42^. (C) All regions of Sin3 subunits with both structural data and self cross-links. Structure mapping and distance measurements were performed using xiVIEW^42^.

### Deleting cross-linking hotspots disrupts Sin3 complex stability

Having observed cross-link hotspots within SIN3A, we reasoned that if a hotspot of cross-links resulted from an important structural interface between SIN3A and the other cross-linked subunits, then deleting regions overlapping these hotspots would result in the loss of binding of the subunits. Therefore, we tested three SIN3A mutants, with either PAH3, part of the HID or PAH4 deleted, for their ability to capture other Sin3 subunits (Fig. 4*A*). The HID 688-829 region appears to be a major interaction interface and cross-links to 5 subunits (SAP30L, SUDS3, BRMS1L, HDAC1 and SAP130—Fig. 4*B*, red lines), whereas the PAH4 region cross-links to 3 subunits (HDAC1, SAP30 and BRMS1L— Fig. 4*B*, green lines) and peptides from only two subunits linked to the PAH3 region (SAP30L and HDAC1—Fig. 4*B*, blue lines) (Fig. 4*B*). Deleting SIN3A HID 688-829 resulted in loss of capture of the subunits cross-linking to this region (or close to this region – BRMS1), as well as loss of capture of HDAC2 and ING1, which are linked to this region via other subunits (HDAC1 and SAP130) and may require these proteins for SIN3A capture (Fig. 4*C*, Supplementary Table 4). In contrast, HID 688-829 removal does not disrupt capture of RBBP7, which cross-links to a distal region at the C terminus of SIN3A. Unlike HID disruption, removal of the PAH3 domain only results in the loss of SAP30L. Although SAP30L cross-links to other regions within SIN3A, to HDAC1, and to BRMS1L, it seems likely that its interaction with PAH3 is required for its stable integration into Sin3 complexes. Disruption of PAH4 had a similar but more modest effect than disruption of the HID suggesting that this region is also involved in stabilizing SIN3A interactions with multiple Sin3subunits. Taken together, the results of Figure 4 confirm that the interprotein cross-links that we have identified map to important regions involved in Sin3 complex stability.

**Figure 4.**
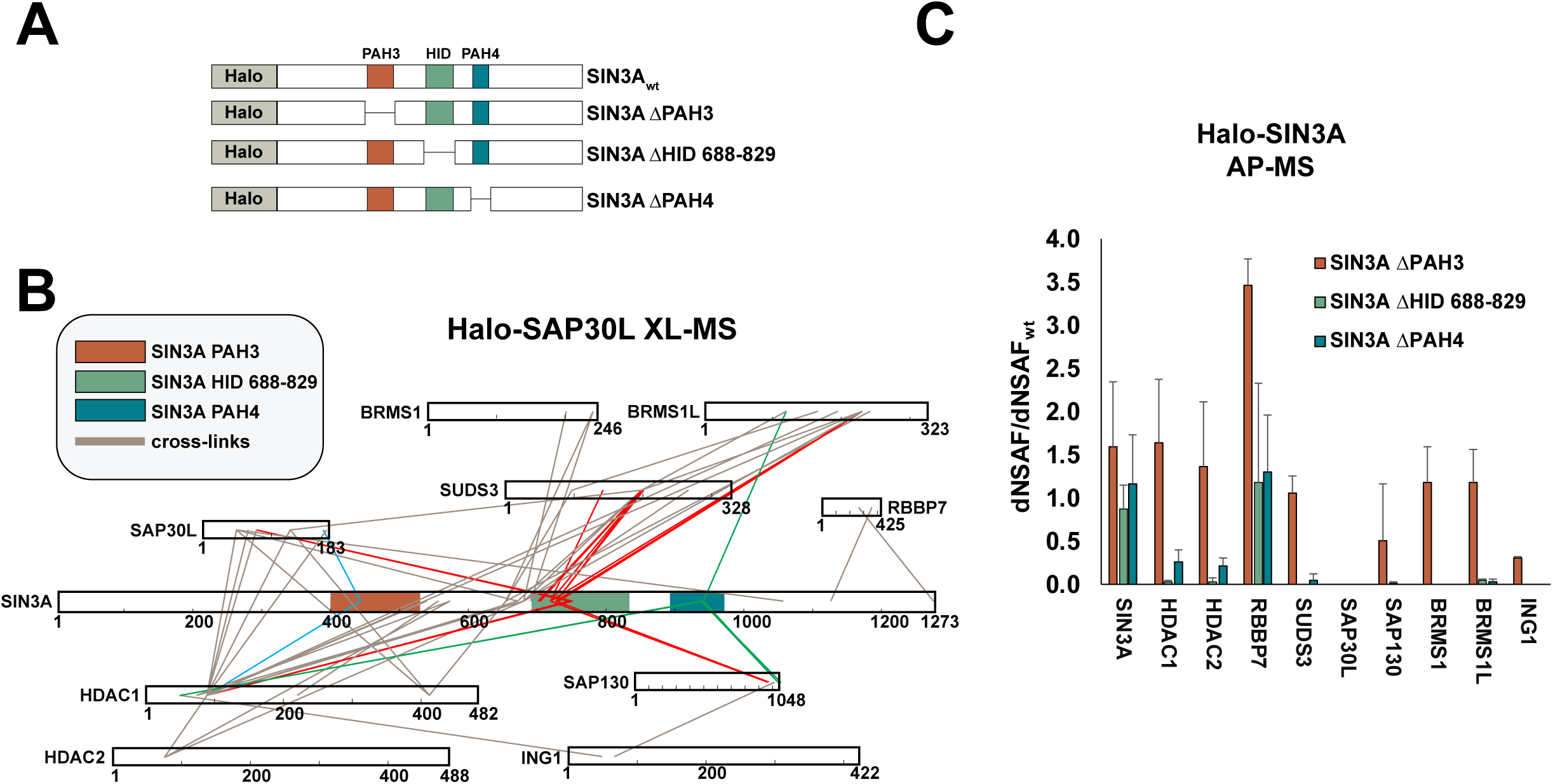
Deletion analysis of SIN3A crosslink hotspots. (A) Regions of SIN3A deleted: Paired Amphipathic Helix 3 (PAH3); Paired Amphipathic Helix 4 (PAH4); and a region 688-829 within the HDAC Interaction Domain (HID). (B) Cross-link map for Sin3 subunit interprotein cross-links. Cross-links to PAH3 (blue), HID 688-829 (red) and to PAH4 (green) are highlighted. (C) Relative abundance of the cross-linked Sin3 subunits shown in (B) copurifying with the SIN3A deletion mutants in AP-MS experiments. Error bars represent standard deviation of 3 biological replicates (Halo-SIN3A ΔPAH3 and Halo-SIN3A ΔHID) or 4 biological replicates (Halo-SIN3A ΔPAH4). Values for each replicate are in Supplementary Table 4.

### Docking SAP30L substructures

Having assessed the validity of our cross-linking data, we next sought to use information from the interprotein cross-links to dock Sin3 structures together. We initially considered how to use the two structures that mapped to SAP30L, 2N1U and 2LD7, which covered most of the N and C terminal halves of SAP30L, respectively. There were 3 cross-links that bridged these structures (shown in blue – Supplementary Fig. 3). As we already had evidence that SAP30L might exist as a homodimer, we did not know whether these cross-links bridge the N and C terminal halves of one SAP30L molecule or alternatively bridge the N terminus of one SAP30L molecule to the C terminus of a second molecule of SAP30L. As the docking of the two structures could be done without resolving which of these possibilities were true, we decided to use these cross-links as restraints when docking SAP30L substructures using the HADDOCK platform^24^ (Supplementary Fig. 3).

### Building a SIN3A/SAP30L/HDAC1 Sin3 complex subassembly guided by cross-linking based restraints

To better understand the 3D architecture of Sin3 complexes, we next used 11 interprotein cross-links to guide docking of the structures mapping to SIN3A, HDAC1, and SAP30L using the HADDOCK platform (Figure 5*A*). We observed several important features in the resulting subassembly.

**Figure 5.**
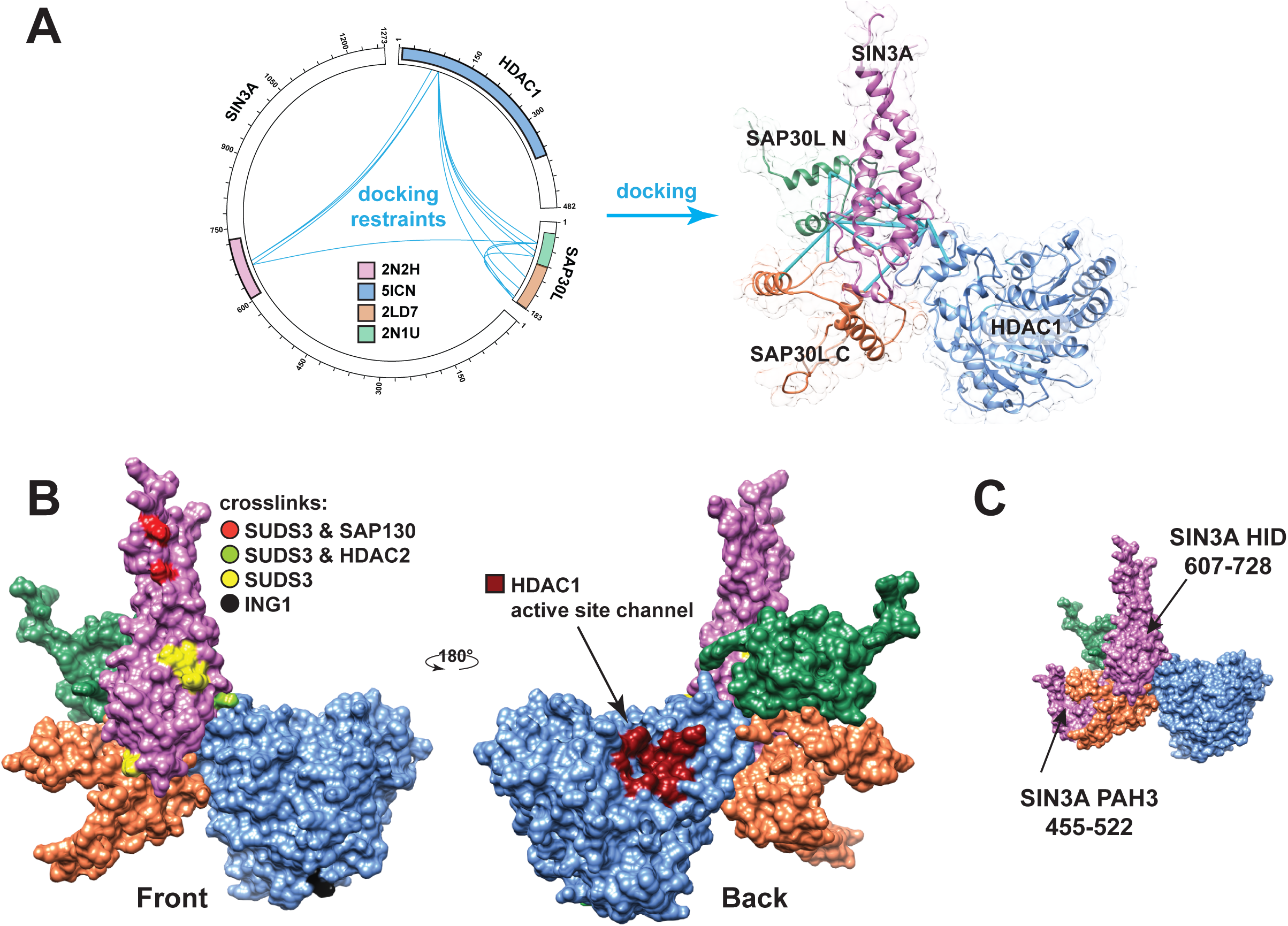
Architecture of the SIN3A/SAP30L/HDAC1 complex. (A) Four Sin3 structures were docked using the HADDOCK platform^44^ guided by docking restraints (Cα–Cα distances < 30 Å) from the indicated cross-links (blue lines). SAP30 structure 2LD7^20^ was mapped to the homologous protein SAP30L using SWISS-MODEL^43^. (B) Structural arrangement of SAP30L, SIN3A and HDAC1, showing the SIN3A residues cross-linked to SUDS3 (yellow), to both SUDS3 and SAP130 (red), to both SUDS3 and HDAC2 (light green), or showing the HDAC1 residue cross-linked to ING1 (black). The position of the HDAC1 active site channel is also shown (dark red). (C) Position of the SIN3A PAH3 domain (previously modeled relative to SAP30 in the structure 2LD7^20^) relative to SAP30L and the SIN3A HID. N.B. although HDAC1 cross-links to the SIN3A PAH3 domain at K439 (lower blue cross-link, Fig. 4*B*), this cross-link does not map to the part of the PAH3 structure shown in Figure 5*C*.

First, after we had docked SAP30L and HDAC1 structures with the SIN3A 607-728 HID platform, there was still access to a large surface on this SIN3A platform onto which other subunits could also assemble (Fig. 5*B*). Indeed, other key Sin3 subunits SUDS3, SAP130 and HDAC2 cross-linked to the remaining exposed front surface of the SIN3A platform (Fig. 5*B*— residues highlighted in yellow, red, and light green).

Second, ING1 cross-links underneath the catalytic subunit HDAC1. Here, ING1 could influence HDAC1 behavior, either by regulating the conformation of HDAC1 and potentially access to the active site, or by helping to position HDAC1 appropriately relative to the histone tail substrate. Some previous evidence is consistent with these possibilities. Smith et al. previously found that treating Sin3/HDAC complexes with the chemotherapeutic drug vorinostat, which enters the active site channel to inhibit HDAC activity, causes ING proteins to dissociate from Sin3/HDAC complexes^28,29^. As we now know that ING1 is proximal to HDAC1, this dissociation might be explained if binding of vorinostat causes an HDAC1 conformational change that precludes binding of ING1/2 or vice versa. In addition, structural studies also provide evidence that ING2, an ING1 homolog, binds H3K4 trimethylated histone tails^30^ and so its proximity to HDAC1 revealed here would be consistent with a model in which ING proteins facilitate favorable positioning of histone tails relative to the HDAC1 active site.

Third, docking both SAP30L N- and C-terminal structures to the SIN3A 667-728 HID platform does not obstruct a likely SAP30L interaction with the SIN3A PAH3 structure. The probable position of part of the SIN3A PAH3 domain (amino acids 455 to 522) relative to SAP30L can be inferred from the structure of SAP30 (a SAP30L homolog) bound to the mSin3A PAH3 domain^20^ (Fig. 5*C*).

## Discussion

A detailed understanding of Sin3 complex architecture facilitates our understanding of how Sin3 complex subunits function in concert to first recruit the HDAC1/2 deacetylases to genomic loci, and then to orient them to coordinate timely deacetylation of histone tails. To capture positional information for Sin3 complex components, we have combined Halo affinity purification with MS cleavable cross-linking techniques to map interface points between or within 13 Sin3 complex subunits.

### The SIN3A HID provides a platform for docking other Sin3 subunits

Cross-linking hotspots on the scaffolding protein SIN3A were centered on the <u>H</u>DAC <u>I</u>nteraction <u>D</u>omain (HID), which is sufficient for transcriptional repression when recruited to promoters in reporter assays^16^. The region SIN3A 607-728 within the HID (previously characterized as the minimal region of mouse Sin3A needed for SUDS3 interaction^21^) forms a platform around which the subunits SUDS3, SAP30/30L, SAP130, HDAC2 and HDAC1 congregate, conceivably to position HDAC1 (and HDAC2) correctly for histone tail deacetylation. These subunits may play subtly different roles in enabling proper HDAC function. SUDS3 forms homodimers and might be involved in tethering two Sin3 complexes together^21^. SAP30L appears to make direct contact with HDAC1 itself and could either guide HDAC1 positioning relative to the SIN3A platform or could influence HDAC1 conformation and hence its activity. SAP30/SAP30L also interact with both core histones and DNA^31^ and Viiri et al. proposed that this interaction helps stabilize Sin3 complexes on nucleosomes. SAP130 has been shown to interact with corepressors and coactivators and likely co-ordinates Sin3 interactions with other coregulatory complexes^19^.

ING1, while not associated with the SIN3A HID platform, cross-links to HDAC1 (Fig. 5*B*). Here, ING1 might help position the HDAC1 correctly relative to histone tail substrate. Indeed, ING proteins do associate with H3K4 trimethylated histone tails^30,32^ and, adjacent to HDAC1, ING1 could offer other acetylated lysines in the histone tail to HDAC1 for deacetylation. Other evidence supports a model in which a direct, controlled interaction between ING proteins and HDAC1 could first direct HDAC1/substrate engagement and then ING protein disengagement. In particular, Sin3/HDAC complexes can lose binding of ING1/2 when the active site channel binds SAHA^28,29^ (vorinostat), an HDAC inhibitor which binds the active site channel in class I HDACs ^33^). Loss of ING protein binding might be explained by inhibitor (or substrate) binding causing HDAC1 conformational changes, which in turn abrogates the HDAC1/ING interaction.

### Distribution of cross-links among Sin3 subunits

The appearance of cross-linking hotspots rather than cross-links evenly distributed among copurified subunits (Fig. 2*C*) could have several explanations. First, we noted that the lowest abundance Sin3 proteins always generated low numbers of cross-links (many proteins cluster near the origin in Fig. 2*D*). Second, the low numbers of cross-links in higher abundance proteins or in regions of these proteins could simply originate from a dearth of lysine residues in these proteins, with “failure to cross-link” not resulting from structural constraints. Although the cross-link deficient RBBP4/7 proteins have the lowest number of lysine residues, this would not fully explain their low number of cross-links. The range of lysine abundancy among all subunits is modest, ranging from ~5-10%, and Sin3 proteins with moderately higher lysine abundances than RBBP4/7 have vastly higher numbers of cross-links/length (compare RBBP4 and SIN3A, Fig. 2*B*). Third, the absence of cross-links might originate from the structural arrangement of Sin3 complexes. Crosslinks might be confined to structured regions rich in alpha helices (there is evidence that β sheets do not yield cross-links^23^), with more dynamic unstructured regions not providing opportunities for self or interprotein cross-linking.

### Novel interfaces

Some cross-links support novel points of contact within and between Sin3 proteins. Self cross-links within SIN3A suggest that regions that are distant in the amino acid chain might be close in Euclidian space. For example, residue 1122 near the SIN3A C-terminus is proximal to residue 747, which resides in the HID. Surprisingly, we found that RBBP7 also cross-linked to the SIN3A C-terminus distal to the HID. An independent interaction between RBBP7 and the C-terminus of SIN3A might explain why RBBP7 is not lost with other subunits when the SIN3A HID 688-829 is deleted (Fig. 4*C*).

### Cross-links with overlapping peptides support dimeric Sin3 complex models

The detection of self cross-links with overlapping peptides suggesting that Sin3 subunit SUDS3 might form homodimers is consistent with previous studies^21,22^. Clark et al. proposed a model to explain SUDS3 dimerization in which two Sin3 complex assemblies operating between adjacent nucleosomes are tethered together by coiled coils in the SUDS3 dimerization domain. Supporting a model in which two Sin3 complexes are bound together to function as a pair, we found self cross-links with overlapping peptides on two other subunits, SAP30L and SIN3A. In addition, ING proteins can homodimerize and ING4 dimers might bind two histone H3 tails either on the same or on different nucleosomes^34^. To conclude, our results are consistent with previous models suggesting that Sin3 complexes might not operate in isolation but instead in pairs, possibly bridging adjacent nucleosomes.

In conclusion, by combining Halo affinity capture with XL-MS using the MS cleavable DSSO cross-linker, we have been able to gain valuable new insight into the relative positioning of Sin3 complex subunits. Our high-confidence cross-link identifications are consistent with existing structural data and with known Sin3 subunit interactions. They highlight novel interfaces between Sin3 subunits and facilitate docking of existing structures, providing new perspectives on Sin3 complex architecture and function.

## Materials and Methods

### Materials

Plasmid FHC11647 and MagneHalo™ affinity beads (#G7281) were purchased from Promega (Madison, WI). 293T cells (ATCC® CRL-11268 ™) were from American Type Culture Collection (Manassas, VA). Salt Active Nuclease (SAN) was from ArcticZymes (Tromso, Norway). AcTEV protease (#12575015) was from Thermo Fisher Scientific (Waltham, MA).

### Cloning Halo-SIN3A wt and deletion mutants for transient expression

Plasmid FHC11647 (Promega, Madison, WI) coding for Halo-SIN3A in pFN21A was altered by site directed mutagenesis (A109V) to code for human SIN3A (Q96ST3/NP_001138830) and to insert a stop codon immediately upstream of the PmeI restriction site at the 3’ end of the ORF. This plasmid was then used as a template, together with the primers listed in Supplementary Data, to clone the SIN3A deletion mutants as follows. First, PCR products were generated corresponding to the portion of the SIN3A ORF 5’ to the deletion site. These N-terminal fragments of SIN3A, flanked by SgfI and KpnI restriction sites were then cloned between the PacI and KpnI sites in plasmid pcDNA5/FRT PacI PmeI previously described^35^. C-terminal fragments of SIN3A downstream of the deletion site flanked by KpnI and PmeI sites were then generated by PCR and inserted between the KpnI and PmeI sites immediately downstream from the N terminal fragments. This resulted in deletion versions of SIN3A with the deleted region replaced by the six base-pair KpnI sequence GGT ACC coding for Gly-Thr.

### Preparation of whole cell lysates from Flp-In™-293 cells stably expressing Halo-SAP30L for XL-MS

A Flp-In™-293 cell line stably expressing Halo-SAP30L expression using a CMV promoter was made essentially as described previously^11^. Approximately 1 × 10^9^ cells were harvested, washed twice in ice cold PBS and frozen at −80°C overnight. Cells were lysed by dounce homogenization in ice-cold lysis buffer containing 20 mM HEPES (pH 7.5), 1.5 mM MgCl_2_, 0.42 M NaCl, 10 mM KCl, 0.2% Triton X-100, 0.5 mM DTT, 0.1 mM benzamidine HCl, 55 μM phenanthroline, 10 μM bestatin, 20 μM leupeptin, 5 μM pepstatin A, 1 mM PMSF, and 500 units SAN (Salt Active Nuclease) and subsequently incubated for 2 hours at 4°C. Lysates were centrifuged at 40, 000 x g at 4°C for 30 minutes and the salt concentration of the resulting supernatants lowered to 0.3 M NaCl by adding an appropriate volume of ice-cold buffer containing 10 mM HEPES (pH 7.5), 1.5 mM MgCl_2_,10 mM KCl, 0.5 mM DTT, 0.1 mM benzamidine HCl, 55 μM phenanthroline, 10 μM bestatin, 20 μM leupeptin, 5 μM pepstatin A, 1 mM PMSF. Lysates were again centrifuged at 40,000 x g at 4°C for 30 minutes and the resulting supernatant harvested for Halo affinity purification of Sin3 complexes for XL-MS experiments.

### Halo purification and cross-linking of complexes from human cells for XL-MS analysis

Lysates prepared from Halo-SAP30L expressing cells were incubated overnight at 4°C with MagneHalo™ magnetic beads prepared from 200 μl bead slurry according to the manufacturer’s instructions. Beads were isolated using a magnetic particle concentrator and washed 4 times in buffer containing 10 mM HEPES (pH 7.5), 1.5 mM MgCl_2_, 0.3 M NaCl, 10 mM KCl, and 0.2% Triton X-100. Bound proteins were eluted by incubating the beads with 200 μl buffer containing 50 mM HEPES (pH 7.5), 0.5 mM EDTA, 1 mM DTT and 30 units AcTEV for at least 2 hours at 4°C. The resulting eluate was recovered, and proteins cross-linked in 5 mM DSSO for 40 minutes at room temperature. The cross-linking reaction was quenched with 50 mM NH_4_HCO_3_ for 15 minutes at room temperature.

### Preparation of whole cell lysates from transiently transfected 293T cells for AP-MS

Approximately 1 × 10^7^ 293T cells were transfected with 7.5 μg of plasmid DNA expressing wt or deletion mutant versions of Halo-SIN3A. Approximately 48 hours after transfection, cells were harvested, washed twice in ice cold PBS, and frozen at −80°C for at least 30 minutes. Cell pellets were resuspended in lysis buffer containing 50 mM Tris•HCl (pH 7.5), 150 mM NaCl, 1% Triton® X-100, 0.1% sodium deoxycholate, 0.1 mM benzamidine HCl, 55 μM phenanthroline, 10 μM bestatin, 20 μM leupeptin, 5 μM pepstatin A and 1 mM PMSF. Lysates were passed through a 26-gauge needle 5-10 times and centrifuged at 21,000 x g for 30 minutes at 4 °C. The resulting supernatant was diluted with 700 μl Tris-buffered saline (25 mM Tris•HCl (pH 7.4), 137 mM NaCl, 2.7 mM KCl).

### Halo purification of complexes from human cells for AP-MS analysis

Lysates prepared from 1 × 10^7^ 293T cells as described above were incubated for 2 hours at 4°C with MagneHalo™ magnetic beads prepared from 100 μl bead slurry according to the manufacturer’s instructions. Beads were washed 4 times in buffer containing 50 mM Tris•HCl (pH 7.4), 137 mM NaCl, 2.7 mM KCl and 0.05% Nonidet® P40. Beads were incubated in elution buffer containing 50 mM Tris•HCl (pH 8.0), 0.5 mM EDTA, 1 mM DTT, and 2 Units AcTEV™ for 2h at 25 °C to elute bound proteins.

### Digestion of proteins for mass spectrometry

Halo purified protein complexes were precipitated by incubation with ice-cold trichloroacetic acid (20% final concentration) overnight at 4°C. Precipitated protein pellets were isolated by centrifugation at 21,000 x g for 30 minutes at 4°C, washed twice in ice-cold acetone, dried, and resuspended in buffer containing 100 mM Tris•HCl (pH 8.5), and 8 M urea. Disulfide bonds were reduced with Tris(2-carboxylethyl)-phosphine hydrochloride. Samples were then treated with chloroacetamide to prevent di-sulfide bond reformation. Denatured proteins were then treated with 0.1 μg Lys-C for 6 hours at 37°C. The urea concentration was reduced to 2 M by adding an appropriate volume of 100 mM Tris•HCl (pH 8.5) and CaCl_2_ added to a final concentration of 2 mM. Proteins were further digested overnight with 0.5 μg trypsin, after which formic acid was added to a final concentration of 5%.

### AP-MS mass spectrometry analysis

For AP-MS experiments, digested samples were loaded onto microcapillary columns containing three phases of chromatography resin (reverse phase, strong cation exchange, reverse phase) and eluted into LTQ mass spectrometers (Thermo Scientific, San Jose, CA) for MudPIT analysis with ten 2-hour chromatography steps^36^. The software package RAWDistiller v. 1.0 was used to convert the resulting .raw files to .ms2 files. The ProLuCID algorithm version 1.3.5^37^ was used to match 36628 human protein sequences (National Center of Biotechnology Information, June 2016 release), 199 common contaminants to MS2 spectra. In addition, the database contained shuffled versions of all sequences for estimating false discovery rates (FDRs). The database was searched for peptides with +57 Da static modifications on cysteine residues (carboxamidomethylation) and with +16 Da dynamic modifications on methionine residues (oxidation). Mass tolerances of 500 ppm (precursor ions) and 500 ppm (fragment ions) were used. Only fully tryptic peptides were considered. The in-house software algorithm Swallow was used in combination with DTASelect^37^ to filter out inaccurate matches and limit spectral, peptide and protein FDRs as described previously^11^. A minimum peptide length of 7 amino acids was established using a DTASelect filter. Proteins that were subsets of others were removed with the parsimony option in Contrast^38^. The in-house software platform NSAF7 was used to calculate dNSAF values using spectral counting^39^. We have reported some of the mass spectrometry data included in this study previously.

Supplementary Table 1 contains a list of all mass spectrometry runs used for this study, spectral and protein FDRs for each run, and details of where each run was first reported. In addition, Supplementary Table 5 contains comprehensive results of the Contrast/NSAF7 analysis of AP-MS data, including peptide-spectrum matches (PSMs). Mass spectrometry raw data has been deposited either to the PeptideAtlas repository^40^ (www.peptideatlas.org), or in the MassIVE repository (http://massive.ucsd.edu). Identifiers for locating the raw data in these repositories are listed in Supplementary Table 1.

### XL-MS mass spectrometry analysis

Cross-linked peptides were resolved for mass spectrometry analysis using a Dionex UltiMate 3000 RSLCnano liquid chromatography system. Peptides were initially loaded from the autosampler onto an Acclaim™ PepMap™ 100 C18 LC Trap Cartridge (0.3 mm inside diameter, 5 mm length) (Thermo Scientific, San Jose, CA) using a loading pump flow rate of 2 μl/minute. Peptides were subsequently resolved for mass spectrometry analysis using an analytical column (50 μm inside diameter, 150 mm length) packed in-house with ReproSil®-Pur C18-aQ 1.9 μm resin (Dr. Masch GmbH, Germany). Chromatography was performed using combinations of buffer A (95% water, 5% acetonitrile, and 0.1% formic acid (v/v/v), pH 2.6), and buffer B (20% water, 80% acetonitrile, and 0.1% formic acid (v/v/v), pH 2.6). The following chromatography steps were performed using a flow rate of 120 nl/minute: (1) 2% B for 20 minutes (column equilibration); (2) a linear gradient from 2% to 10% B over 10 minutes; (3) a linear gradient from 10% to 40% B over either 120 minutes or 240 minutes; (4) a linear gradient from 40% to 95% B over 5 minutes; (5) 95% B for 14 minutes (column wash); (6) a linear gradient from 95% B to 2% B over 1 minute; (7) 2% B for 10 minutes (column re-equilibration).

Eluted peptides were analyzed by mass spectrometry using an Orbitrap Fusion™ Lumos™ mass spectrometer (Thermo Scientific, San Jose, CA). An MS3 based method was used for identification of DSSO cross-linked peptides as follows: Full MS scans were performed using the Orbitrap mass analyzer (60,000 m/z resolution, 1.6 m/z isolation window, and 375-1500 m/z scan range); The top 3 peptides identified with charge state 4 to 8 were selected for MS2 fragmentation (20% CID energy) and subsequent detection with the Orbitrap mass analyzer (30,000 m/z resolution and a dynamic exclusion time of 40 s); Pairs of MS2 fragments with a mass difference of 31.9720 (20 ppm mass tolerance) were selected for MS3 fragmentation (CID energy 35%) and detection using the Linear Ion Trap mass analyzer (rapid scan, 3 m/z isolation window, maximum ion injection time 200 ms); Each MS2 scan was followed by a maximum of 4 MS3 scans.

### Identification of DSSO Cross-Linked Peptides

Proteome Discoverer 2.2 with the add on XlinkX cross-linking nodes^41^ (Thermo Scientific, San Jose, CA) was used to identify cross-linked peptides from .raw files from three experiments as follows. For each .raw file, the Xlinkx Detect processing node was used to identify MS2 fragmentation scans with reporter ions characteristic of DSSO crosslinked peptides using DSSO lysine crosslink modifications of 158.00376 Da (monoisotopic mass) and 158.17636 Da (average mass), with cleaved DSSO lysine crosslink modifications of 54.01056 Da (alkene, monoisotopic mass), 54.04749 Da (alkene, average mass), 85.98264 Da (thiol, monoisotopic mass), and 86.11358 Da (thiol, average mass). Subsequently, a version of the database used for AP-MS searches but without shuffled sequences was searched using either the Xlinkx Search node (fragmentation scans with cross-link reporter ions) or with the Sequest HT node (scans without cross-link reporter ions). Both search strategies searched for peptides with 57.021 Da fixed modifications on cysteine residues (carbamidomethylation) and 15.995 Da variable modifications on methionine residues (oxidation). In addition, the Sequest HT node searched for the variable lysine modifications 176.014 Da (water quenched DSSO monoadducts) and 279.078 Da (Tris quenched DSSO monoadducts). For Sequest HT node searches, a precursor ion mass tolerance of 10 ppm and a fragment ion mass tolerance of 0.6 Da were used; for Xlinkx Search node searches, a precursor ion mass tolerance of 10 ppm and fragment ion mass tolerances of 20 ppm (FTMS) or 0.5 Da (ITMS) were used. The maximum number of equal dynamic modifications was 3 (Sequest HT searches). The protein FDR was set at 0.01 using the Xlinkx Validator node for Xlinkx searches, and the target FDR (Strict) was set at 0.01 using the Percolator node for Sequest HT searches. Supplementary Table 6 contains comprehensive details of crosslink identifications, including crosslink-spectrum matches (CSMs). Raw data and Proteome Discoverer results files for XL-MS experiments has been deposited in the MassIVE repository with the identifier MSV000084311 (see Supplementary Table 1).

### Downstream analysis of cross-linking data

A summary of the high-confidence cross-links identified using Proteome Discoverer 2.2 that were used for further analysis is presented in Supplementary Table 3. The xiView platform was used for cross-link visualization, mapping cross-links to PDB structures and calculating distances between alpha carbon atoms^42^. The SWISS-MODEL platform^43^ was used for homology-modeling the PDB files corresponding to human SIN3A (using PDB 2N2H corresponding to mSin3A) and to the C-terminus of SAP30L (using PDB 2LD7 corresponding to human SAP30) used for protein docking in Figure 5. Protein structures were docked using the HADDOCK server^44^ using default settings together with the cross-linking restraints indicated in Figure 5. Protein structures were processed for visualization with Chimera^45^.

### Data Availability Statement

The mass spectrometry datasets generated for this study (used for the analyses in Fig. 2 to Fig. 5) are available from the Massive data repository (https://massive.ucsd.edu) using the identifiers listed in Supplementary Table 1.

### Code Availability

Code for the software RAWDistiller v. 1.0 and NSAF7 is available on request.

## Acknowledgements

Research reported in this publication was supported by the Stowers Institute for Medical Research and the National Institute of General Medical Sciences of the National Institutes of Health under Award Number RO1GM112639 to MPW and F32GM122215 to MKA. The content is solely the responsibility of the authors and does not necessarily represent the official views of the National Institutes of Health.

Original data underlying this manuscript can be accessed from the Stowers Original Data Repository at http://www.stowers.org/research/publications/LIBPB-1465.

## Author Contributions

C.A.S.B., Y.Z., and M.P.W. wrote the manuscript and designed the project. All authors reviewed and commented on the manuscript. L.F. and Z.W. contributed data analysis tools. C.A.S.B. and S.M.I. constructed plasmids. S.M., J.L.T., M.K.A. and C.A.S.B. performed AP-MS analyses Y.Z. and Y.H. performed XL-MS analyses. M.P.W. supervised the project.

## References

1. Smialowski, P. & Wong, P. Protein Crystallizability. in Data Mining Techniques for the Life Sciences (eds. Carugo, O. & Eisenhaber, F.) 341–370 (Springer New York, 2016). doi:10.1007/978-1-4939-3572-7_17

2. Frueh, D. P., Goodrich, A. C., Mishra, S. H. & Nichols, S. R. NMR methods for structural studies of large monomeric and multimeric proteins. Curr. Opin. Struct. Biol. 23, 734–739 (2013).

3. Leitner, A., Faini, M., Stengel, F. & Aebersold, R. Crosslinking and Mass Spectrometry: An Integrated Technology to Understand the Structure and Function of Molecular Machines. Trends Biochem. Sci. 41, 20–32 (2016).

4. Rappsilber, J., Siniossoglou, S., Hurt, E. C. & Mann, M. A generic strategy to analyze the spatial organization of multi-protein complexes by cross-linking and mass spectrometry. Anal. Chem. 72, 267–275 (2000).

5. Kao, A. et al. Development of a novel cross-linking strategy for fast and accurate identification of cross-linked peptides of protein complexes. Mol. Cell. Proteomics 10, 1–17 (2011).

6. Wang, X. et al. Molecular details underlying dynamic structures and regulation of the human 26S proteasome. Mol. Cell. Proteomics 16, 840–854 (2017).

7. Los, G. V et al. HaloTag: a novel protein labeling technology for cell imaging and protein analysis. ACS Chem. Biol. 3, 373–382 (2008).

8. Kelly, R. D. W. & Cowley, S. M. The physiological roles of histone deacetylase (HDAC) 1 and 2: complex co-stars with multiple leading parts. Biochem. Soc. Trans. 41, 741–9 (2013).

9. Zhang, Y. et al. Analysis of the NuRD subunits reveals a histone deacetylase core complex and a connection with DNA methylation. Genes Dev. 13, 1924–1935 (1999).

10. Lee, M. G., Wynder, C., Cooch, N. & Shiekhattar, R. An essential role for CoREST in nucleosomal histone 3 lysine 4 demethylation. Nature 437, 432–435 (2005).

11. Banks, C. A. S. et al. A Structured Workflow for Mapping Human Sin3 Histone Deacetylase Complex Interactions Using Halo-MudPIT Affinity-Purification Mass Spectrometry. Mol. Cell. Proteomics 17, 1432–1447 (2018).

12. Kandoth, C. et al. Mutational landscape and significance across 12 major cancer types. Nature 502, 333–339 (2013).

13. Kwon, Y. J. et al. Selective inhibition of SIN3 corepressor with avermectins as a novel therapeutic strategy in triple-negative breast cancer. Mol. Cancer Ther. 14, 1824–1836 (2015).

14. Rielland, M. et al. Senescence-associated SIN3B promotes inflammation and pancreatic cancer progression. J. Clin. Invest. 124, 2125–2135 (2014).

15. Marks, P. a & Breslow, R. Dimethyl sulfoxide to vorinostat: development of this histone deacetylase inhibitor as an anticancer drug. Nat. Biotechnol. 25, 84–90 (2007).

16. Laherty, C. D. et al. Histone deacetylases associated with the mSin3 corepressor mediate mad transcriptional repression. Cell 89, 349–56 (1997).

17. Zhang, Y. et al. SAP30, a novel protein conserved between human and yeast, is a component of a histone deacetylase complex. Mol. Cell 1, 1021–1031 (1998).

18. Alland, L. et al. Identification of mammalian Sds3 as an integral component of the Sin3/histone deacetylase corepressor complex. Mol. Cell. Biol. 22, 2743–50 (2002).

19. Fleischer, T. C., Yun, U. J. & Ayer, D. E. Identification and Characterization of Three New Components of the mSin3A Corepressor Complex. 23, 3456–3467 (2003).

20. Xie, T. et al. Structure of the 30-kDa Sin3-associated protein (SAP30) in complex with the mammalian Sin3A corepressor and its role in nucleic acid binding. J. Biol. Chem. 286, 27814–24 (2011).

21. Clark, M. D. et al. Structural insights into the assembly of the histone deacetylase-associated Sin3L/Rpd3L corepressor complex. Proc. Natl. Acad. Sci. U. S. A. 112, E3669–78 (2015).

22. Clark, M. D., Zhang, Y. & Radhakrishnan, I. Solution NMR Studies of an Alternative Mode of Sin3 Engagement by the Sds3 Subunit in the Histone Deacetylase-Associated Sin3L/Rpd3L Corepressor Complex. J. Mol. Biol. 427, 3817–3823 (2015).

23. Schneider, M., Belsom, A., Rappsilber, J. & Brock, O. Blind testing of cross-linking/mass spectrometry hybrid methods in CASP11. Proteins Struct. Funct. Bioinforma. 84, 152–163 (2016).

24. Merkley, E. D. et al. Distance restraints from crosslinking mass spectrometry: Mining a molecular dynamics simulation database to evaluate lysine-lysine distances. Protein Sci. 23, 747–759 (2014).

25. Laitaoja, M. et al. Redox-dependent disulfide bond formation in SAP30L corepressor protein: Implications for structure and function. Protein Sci. 25, 572–586 (2016).

26. Watson, P. J. et al. Insights into the activation mechanism of class I HDAC complexes by inositol phosphates. Nat. Commun. 7, 1–13 (2016).

27. Murzina, N. V. et al. Structural Basis for the Recognition of Histone H4 by the Histone-Chaperone RbAp46. Structure 16, 1077–1085 (2008).

28. Smith, K. T., Martin-Brown, S. a., Florens, L., Washburn, M. P. & Workman, J. L. Deacetylase Inhibitors Dissociate the Histone-Targeting ING2 Subunit from the Sin3 Complex. Chem. Biol. 17, 65–74 (2010).

29. Sardiu, M. E. et al. Suberoylanilide hydroxamic acid (SAHA)-induced dynamics of a human histone deacetylase protein interaction network. Mol. Cell. Proteomics 13, 3114–3125 (2014).

30. Peña, P. V. et al. Molecular mechanism of histone H3K4me3 recognition by plant homeodomain of ING2. Nature 442, 100–103 (2006).

31. Viiri, K. M. et al. DNA-binding and -bending activities of SAP30L and SAP30 are mediated by a zinc-dependent module and monophosphoinositides. Mol. Cell. Biol. 29, 342–356 (2009).

32. Shi, X. et al. ING2 PHD domain links histone H3 lysine 4 methylation to active gene repression. Nature 442, 96–99 (2006).

33. Lauffer, B. E. L. et al. Histone deacetylase (HDAC) inhibitor kinetic rate constants correlate with cellular histone acetylation but not transcription and cell viability. J. Biol. Chem. 288, 26926–26943 (2013).

34. Culurgioni, S. et al. Crystal structure of inhibitor of growth 4 (ING4) dimerization domain reveals functional organization of ING family of chromatin-binding proteins. J. Biol. Chem. 287, 10876–10884 (2012).

35. Banks, C. A. S. et al. Controlling for gene expression changes in transcription factor protein networks. Mol. Cell. Proteomics 13, 1510–22 (2014).

36. Banks, C., Kong, S. & Washburn, M. Affinity purification of protein complexes for analysis by multidimensional protein identification technology. Protein Expr. Purif. 86, 105–119 (2012).

37. Xu, T. et al. ProLuCID: An improved SEQUEST-like algorithm with enhanced sensitivity and specificity. J. Proteomics 129, 16–24 (2015).

38. Tabb, D. L., McDonald, W. H. & Yates III, J. R. DTASelect and Contrast: tools for assembling and comparing protein identifications from shotgun proteomics. J. Proteome Res. 1, 21–6 (2002).

39. Zhang, Y., Wen, Z., Washburn, M. P. & Florens, L. Refinements to label free proteome quantitation: how to deal with peptides shared by multiple proteins. Anal. Chem. 82, 2272–81 (2010).

40. Desiere, F. et al. The PeptideAtlas project. Nucleic Acids Res. 34, D655–8 (2006).

41. Liu, F., Lössl, P., Scheltema, R., Viner, R. & Heck, A. J. R. Optimized fragmentation schemes and data analysis strategies for proteome-wide cross-link identification. Nat. Commun. 8, (2017).

42. Graham, M. J., Combe, C., Kolbowski, L. & Rappsilber, J. xiView: A common platform for the downstream analysis of Crosslinking Mass Spectrometry data. bioRxiv 1–5 (2019). doi:10.1101/561829

43. Waterhouse, A. et al. SWISS-MODEL: Homology modelling of protein structures and complexes. Nucleic Acids Res. 46, W296–W303 (2018).

44. De Vries, S. J., Van Dijk, M. & Bonvin, A. M. J. J. The HADDOCK web server for data-driven biomolecular docking. Nat. Protoc. 5, 883–897 (2010).

45. Pettersen, E. F. et al. UCSF Chimera - A visualization system for exploratory research and analysis. J. Comput. Chem. 25, 1605–1612 (2004).

