## Supplementary Information and Figures for "Integrative Modeling of a Sin3/HDAC Complex Sub-structure"

1. **Primers used to construct vectors expressing Halo tagged SIN3A mutants in pcDNA5/FRT PacI PmeI (restriction sites used in red)**

| | |
|---|---|
| SIN3A SgfI F | 5'- CAG GCG ATC GCC ATG AAG CGG CGT TTG GAT GAC C- 3' |
| SIN3A PmeI R | 5'- CAG GTT TAA ACT TAA GGG GCT TTG AAT ACT GTG CCG TAT TTG - 3' |
| SIN3A ΔPAH3 F | 5'- CAT GGT ACC GAG TCT GTA CAT CTG GAA ACT TAT CCA - 3' |
| SIN3A ΔPAH3 R | 5'- CAT GGT ACC CTC AGC AGT TGT TTT GCT TAA AAG C - 3' |
| SIN3A ΔHID F | 5'- CAT GGT ACC GAT CTC TCA GAT GTG GAG GAA GAG GAA - 3' |
| SIN3A ΔHID R | 5'- CAT GGT ACC ATT CTT TCT CAG ACC ATC AAT GAT G - 3' |
| SIN3A ΔPAH4 F | 5'- CAT GGT ACC AGC CTG CTG GAT GGC AAC ATA GAC TCA - 3' |
| SIN3A ΔPAH4 R | 5'- CAT GGT ACC GAC ATA GAA GAG GTT GTA TAC TTC ATC CA- 3' |

**Supplementary Figure 1**

Relative abundance of Sin3 complex subunits copurifying with the Halo-SAP30L subunit determined by AP-MS. Values for each subunit are 1000 x mean dNSAF (data and experimental details previously published in Banks et al. 2018[1]).

**Supplementary Figure 2**

Halo-SAP30L purified complexes treated with or without DSSO cross-linking. Samples purified from Flp-In™-293 cells stably expressing Halo-SAP30L as described in methods were treated with or without 5 mM DSSO for 40 minutes at room temperature, boiled in sample buffer and separated by SDS-PAGE. Proteins were visualized by silver staining.

**Supplementary Figure 3**

Docking structures mapping to SAP30L. SAP30 structure 2LD7 was mapped to the homologous protein SAP30L using SWISS-MODEL. A cross-link with overlapping peptides (red) is consistent with a SAP30L homodimer. The self cross-links shown in blue were then used as docking restraints to assemble the structures using the HADDOCK platform, leaving unresolved the question as to whether the docked structures represent regions of the same SAP30L molecule or two different SAP30L molecules.

**Supplementary Tables:**

**Supplementary Table 1:** Mass spectrometry runs used in this study.

**Supplementary Table 2:** Previously characterized Sin3 subunit interactions.

**Supplementary Table 3:** Cross-links identified in DSSO treated SAP30L purifications

**Supplementary Table 4:** Quantitative analysis of Sin3 subunits and Sin3 associated transcription factors identified in Halo-SIN3A wt and deletion mutant purifications by AP-MS.

**Supplementary Table 5:** Comprehensive results of Contrast/NSAF7 analysis of AP-MS data

**Supplementary Table 6:** Comprehensive details of crosslink identifications, including crosslink-spectrum matches (CSMs)

**Reference used in Supplementary Information**

1. Banks, C. A. S. *et al.* A Structured Workflow for Mapping Human Sin3 Histone Deacetylase Complex Interactions Using Halo-MudPIT Affinity-Purification Mass Spectrometry. *Mol. Cell. Proteomics* **17**, 1432–1447 (2018).

### Supplementary Figure 1

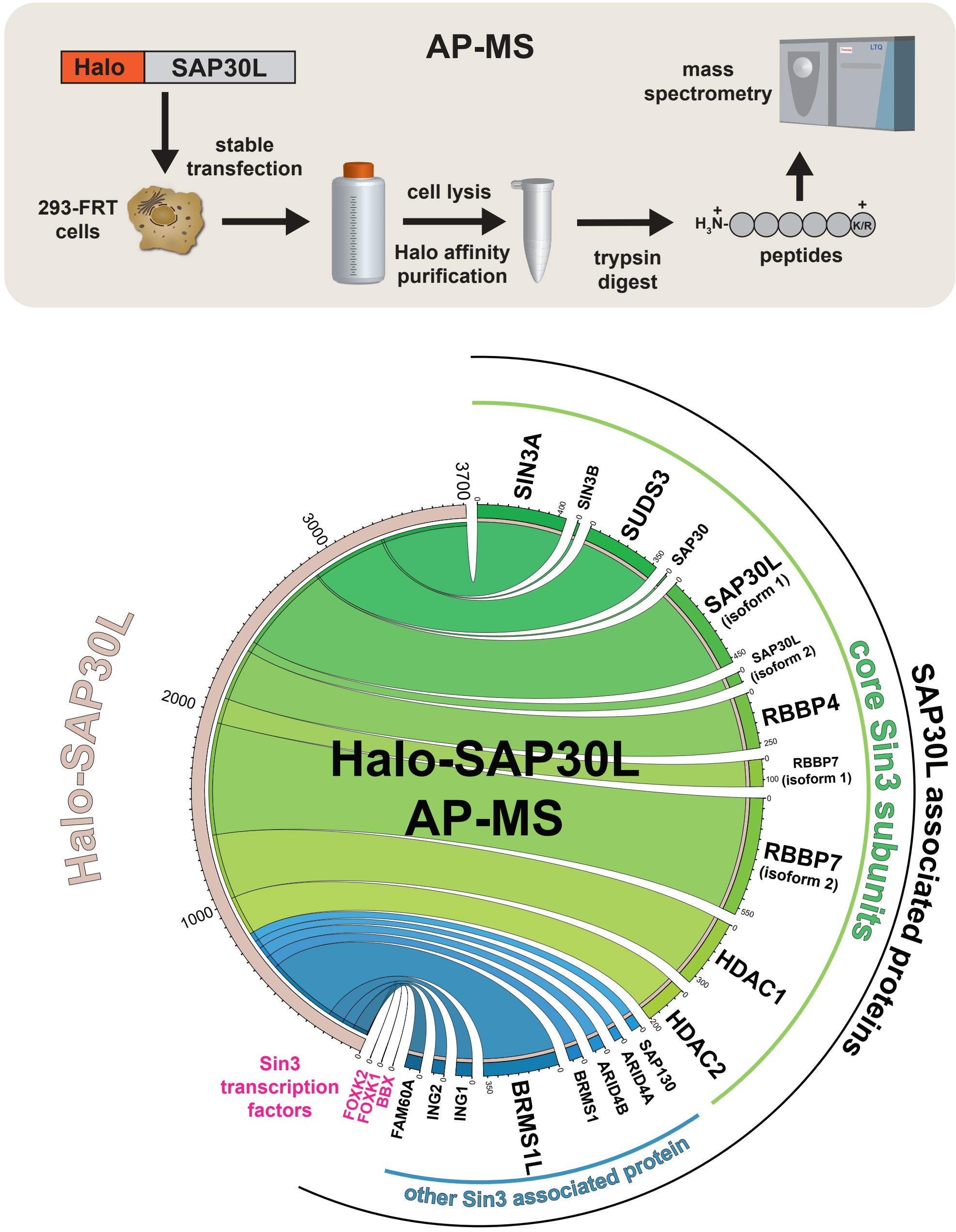


Relative abundance of Sin3 complex subunits copurifying with the Halo-SAP30L subunit determined by AP-MS. Values for each subunit are 1000 x mean dNSAF (data and experimental details previously published in Banks et al. 2018).

**Supplementary Figure 2**

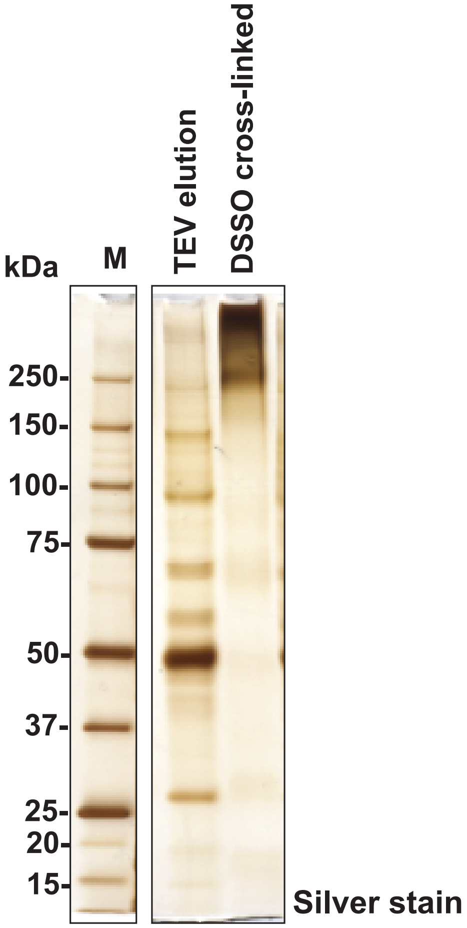


**Halo-SAP30L purified complexes treated with or without DSSO cross-linking.**
Samples purified from Flp-In™-293 cells stably expressing Halo-SAP30L as described in methods were treated with or without 5 mM DSSO for 40 minutes at room temperature, boiled in sample buffer and separated by SDS-PAGE. Proteins were vizualized by silver staining.

**Supplementary Figure 3**

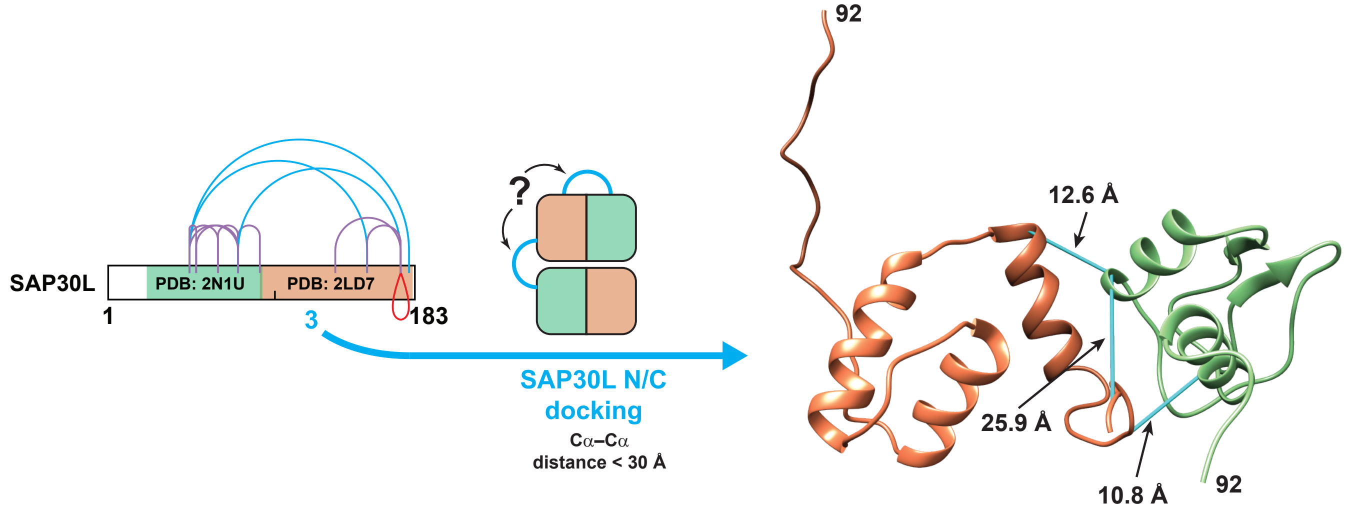


**Docking structures mapping to SAP30L.**
SAP30 structure 2LD7 was mapped to the homologous protein SAP30L using SWISS-MODEL. A cross-link with overlapping peptides (red) is consistent with a SAP30L homodimer. The self cross-links shown in blue were then used as docking restraints to assemble the structures using the HADDOCK platform, leaving unresolved the question as to whether the docked structures represent regions of the same SAP30L molecule or two different SAP30L molecules.
